# Structural basis of INTAC-regulated transcription

**DOI:** 10.1101/2021.11.29.470345

**Authors:** Hai Zheng, Qianwei Jin, Yilun Qi, Weida Liu, Yulei Ren, Xinxin Wang, Fei Chen, Jingdong Cheng, Xizi Chen, Yanhui Xu

## Abstract

For the majority of expressed eukaryotic genes, RNA polymerase II (Pol II) forms a paused elongation complex (PEC) and undergoes promoter-proximal pausing downstream of the transcription start site ^1–3^. The polymerase either proceeds into productive elongation or undergoes promoter-proximal premature transcription termination ^4–6^. It remains incompletely understood how transcription is regulated at this stage. Here, we determined the structure of PEC bound to INTAC, an Integrator-containing PP2A complex ^7^, at near-atomic resolution. The structure shows that INTAC partially wraps around PEC through multiple contacts, permitting the memetic nascent RNA to run into substrate-entry tunnel of the endonuclease subunit INTS11 of INTAC for cleavage. Pol II C-terminal domain (CTD) winds over INTAC backbone module through multiple anchors and is suspended above the phosphatase of INTAC for dephosphorylation. Biochemical analysis shows that INTAC-PEC association requires unphosphorylated CTD and could tolerate CTD phosphorylation, suggesting an INTAC-mediated persistent CTD dephosphorylation followed by reinforcement of the INTAC-PEC complex. Our study reveals how INTAC binds PEC and orchestrates RNA cleavage and CTD dephosphorylation, two critical events in generating premature transcription termination.

## Introduction

Eukaryotic transcription by RNA polymerase II (Pol II) is a strictly regulated process that involves the interplay of numerous factors ^4,5^. Promoter-proximal pausing is a regulatory mechanism that connects transcription initiation and productive elongation in metazoan ^3,6^. It typically occurs at 20–200 base pairs downstream of the transcription start site (TSS) and can be observed at the majority of expressed genes ^1,2^. Pol II forms a paused elongation complex (PEC) through binding of two factors: the 5,6-dichloro-1-β-d-ribofuranosylbenzimidazole (DRB) sensitivity-inducing factor (DSIF), consisting of subunits SPT4 and SPT5, and the negative elongation factor (NELF), consisting of the four subunits NELF-A, -B, -C/D and -E ^8–11^. Following the duration of pausing, the polymerase either proceeds into productive elongation or undergoes promoter-proximal premature transcription termination (PTT) ^12^, which plays a decisive role in determining transcriptional outputs. Nonetheless, it remains unclear how the fate of PEC is governed mechanistically.

In contrast to the well-characterized pause release and productive elongation ^4,5^, the mechanism of PTT remains largely unknown. Emerging evidence implies that metazoan-specific Integrator complex involves this process. Integrator complex ^13,14^ functions as an RNA endonuclease to cleave different classes of RNAs ^15–19^. More recent studies discovered that Integrator is enriched in the proximity of RNA promoters ^20^ and can associate with paused Pol II bound by DSIF and NELF ^21,22^ to trigger PTT and repress gene activity ^18,23–28^. We have recently found that Integrator associates with protein phosphatase 2A core enzyme (PP2A-AC) and dephosphorylates the C-terminal domain (CTD) of Pol II and determined the structure of Integrator-containing PP2A-AC (termed INTAC), showing how the RNA nuclease and protein phosphatase are organized in the INTAC complex ^7^. In addition, Integrator-bound PP2A dephosphorylates Pol II CTD and Spt5 to prevent the transition to productive elongation ^7,24,29^. Despite these studies, it remains elusive how INTAC, especially its two catalytic modules, is structurally organized and functionally coordinated in the context of PEC and how INTAC works with PEC in PTT.

### Structure determination of INTAC-PEC complex

To investigate the mechanistic implications of INTAC in promoter-proximal pausing, we purified the human INTAC ^7^, NELF, DSIF, and pig (S. scrofa) Pol II, which has 99.9% sequence identity to human Pol II except for four amino acids (Extended Data Fig. 1, a, b). The INTAC-PEC complex was assembled by adding INTAC, NELF, and DSIF to an elongation complex that was pre-assembled by mixing Pol II and a DNA-RNA hybrid ^30,31^ (Methods). The assembled INTAC-PEC complex was subjected to gradient fixation (GraFix), followed by cryo-electron microscopy (cryo-EM) single particle reconstruction (Extended Data Fig. 2). The cryo-EM map was refined to 4.5 Å resolution and the maps of subcomplexes were improved to near-atomic (3.8 Å to 4.1 Å) resolution by focused refinement. Structural model was built by fitting previously determined structures of INTAC ^7^ and PEC ^30^ into the cryo-EM maps followed by manual adjustment (Extended Data Fig. 3, Extended Data Table 1).

### Overall structure of INTAC-PEC complex

The INTAC-PEC complex structure reveals a compact fold with approximate dimensions of ~280× 270× 260 Å^3^ (Fig. 1, Extended Data Fig. 4, Supplementary Video 1). As observed in the apo PEC structure ^30^, NELF and DSIF wrap around the central Pol II, generating a compact globular fold. The PEC complex sits above the main body of INTAC through interface-I/-II/-V and is further stabilized by two INTAC protrusions on opposite sides at interface-III/-IV, resembling a ball (PEC) in a bucket (INTAC). Consistent with the modular organization of the apo INTAC complex ^7^, the shoulder and backbone modules of INTAC within INTAC-PEC generate a central cruciform scaffold with the phosphatase and endonuclease modules flanking the opposite sides. A previously undetected tail module extends out of the bottom of the backbone module and folds back to bind Pol II. Three putative Pol II CTD fragments wind on the surface of INTAC backbone module and an additional fragments is suspended above the catalytic pocket of the phosphatase subunit PP2A-C, indicative of a path favorable for INTAC-mediated CTD association and dephosphorylation (Supplementary Video 2).

**Fig. 1.**
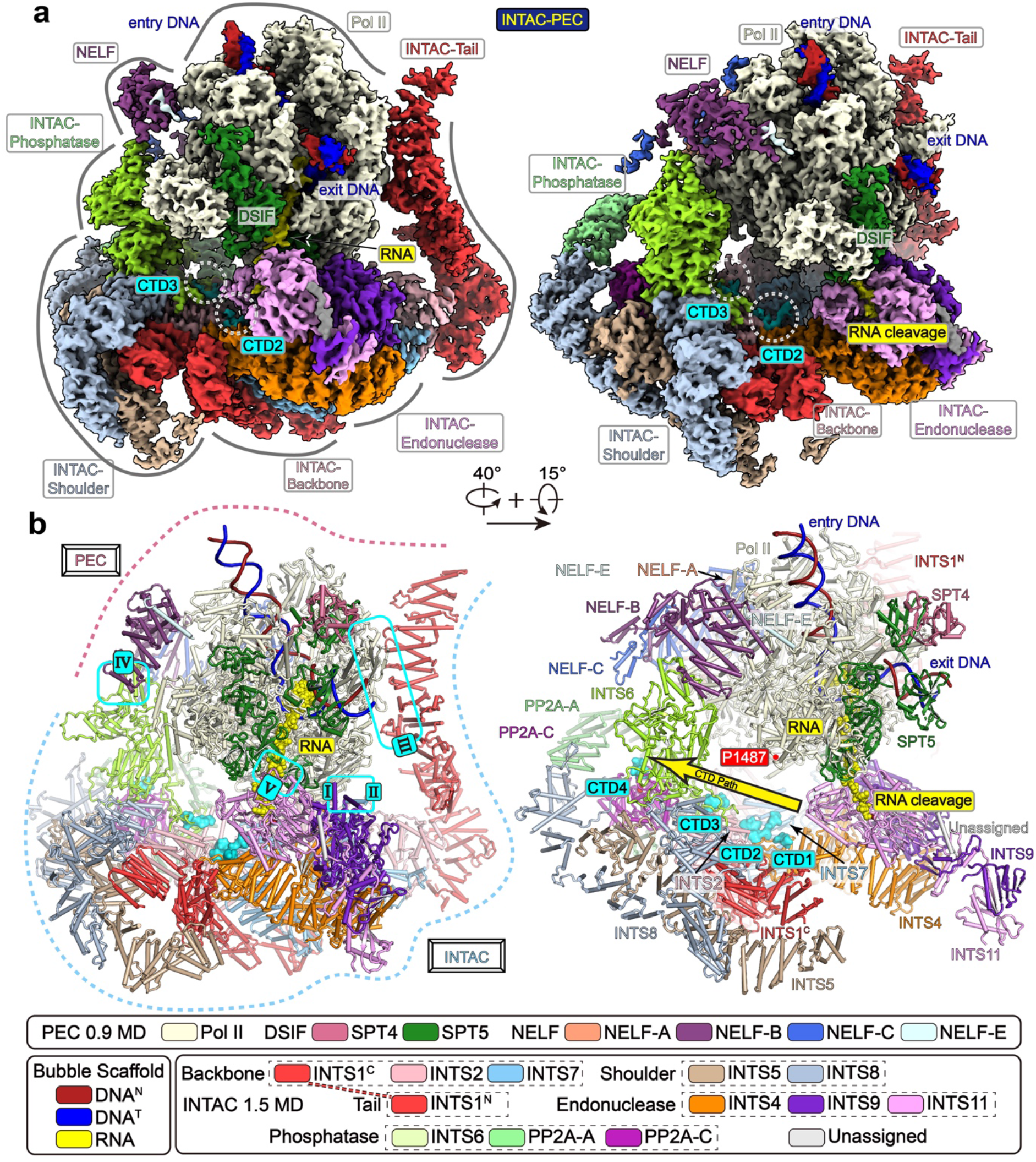
Structure of INTAC-PEC complex. **a, b,** Composite cryo-EM map (**a**) and structural model (**b**) of INTAC-PEC shown in two different views. Five INTAC-PEC interfaces are indicated and four putative Pol II CTD fragments are shown in surface representation. The same color scheme was used in all of the figures if not otherwise specified. Putative CTD-binding path on INTAC and RNA cleavage site in INTS11 are highlighted.

The DNA duplex is opened in the catalytic cavity with the template strand forming a one-turn DNA-RNA hybrid, and reanneals into a duplex that protrudes out of Pol II through the exit tunnel (Fig. 1b, Extended Data Fig. 4). Both exit and entry DNAs point away from INTAC, suggesting that INTAC generates no clash with DNA and that INTAC does not directly affect transcription elongation. The memetic nascent RNA runs out through the RNA exit tunnel of Pol II, is brought into proximity of INTAC by Pol II-SPT5-INTS11 interactions at interface-V, and runs into the RNA entry tunnel of the endonuclease subunit INTS11. The last visible RNA nucleotide is suspended above the active site of INTS11. The structure reveals INTAC-PEC organization that favors RNA cleavage and Pol II CTD dephosphorylation.

### The interfaces between INTAC and Pol II-NELF

INTAC makes three direct contacts with Pol II (Fig. 2, a to d, Extended Data Fig. 5). At the interface-I, the C-terminal α-helix of RPB11 contacts the helical repeat 1 of INTS2 (INTS2^HR1^). At the interface-II, the C-terminal helix of INTS7 binds the domain2 of RPB3 and the C-terminal end of INTS4 bridges the contact between INTS9 and the zinc loop of RPB3. The N-terminal HEAT (huntingtin, elongation factor 3, protein phosphatase 2A and TOR1) domain of INTS1 adopts an arch-shaped fold and forms the tail module, which caps the exposed end of the scaffold module and flanks away from the core INTAC. The two RPB2 external domains of Pol II contact the tail on the convex ridge at the interface-III. This tail module was not observed in the apo INTAC structure (Extended Data Fig. 4a), suggesting a PEC-mediated tail stabilization and a potential function of tail in recruitment of PEC.

**Fig. 2.**
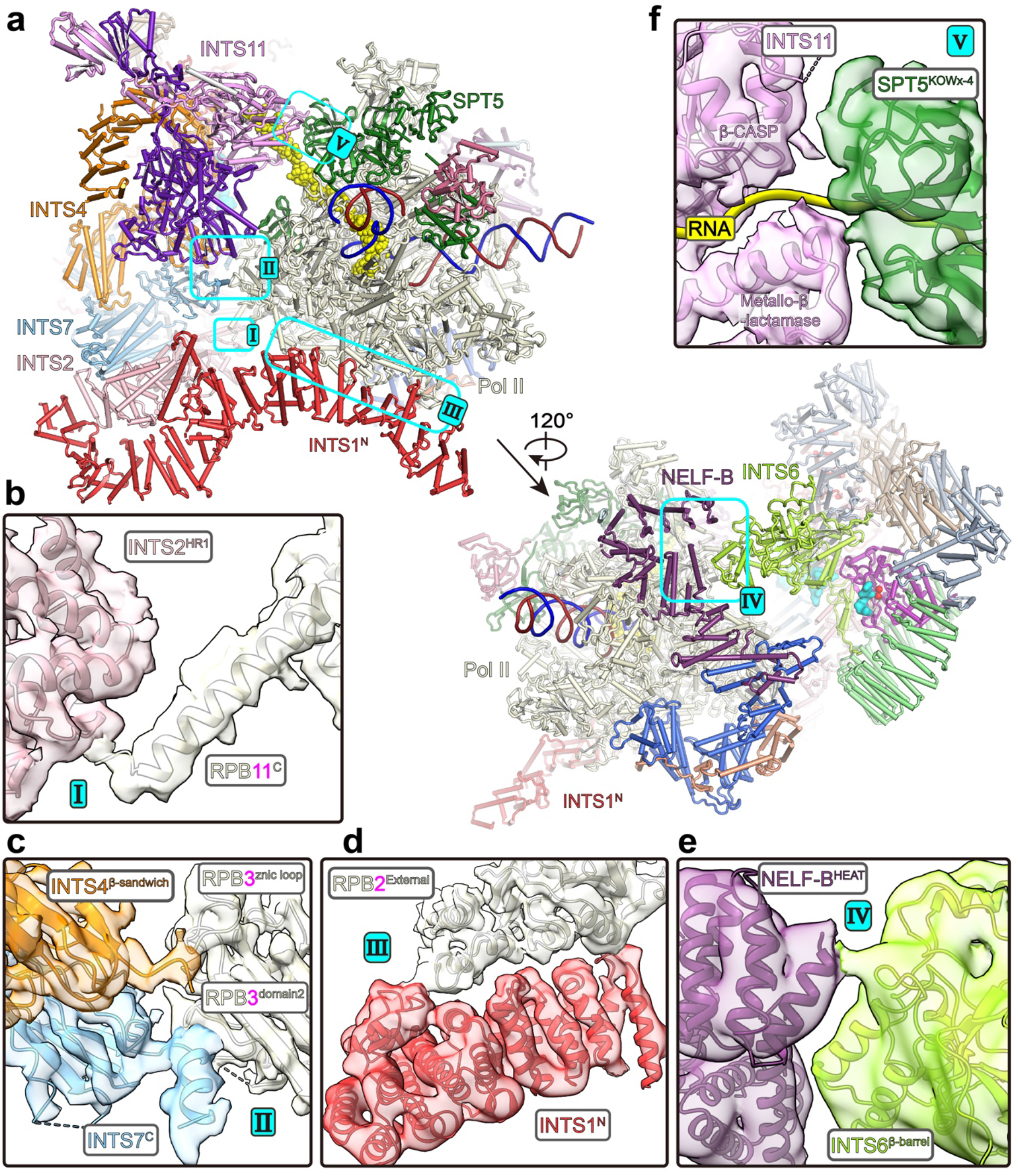
Interactions between INTAC and PEC. **a,** Overall structure of INTAC-PEC with the five inter-complex contacts highlighted. **b-f,** Close-up views of the interactions with cryo-EM maps shown in transparent surface and structural models shown in cartoon.

INTS6 bridges the phosphatase of INTAC and the NELF-B-NELF-E lobe ^30^ of PEC (Fig. 2, a, e, Extended Data Fig. 5). At the interface-IV, the exposed end of the INTS6 β-barrel domain contacts the HEAT domain of NELF-B, consistent with known interaction between Integrator and NELF ^21,22^. In addition, INTS6 ^7^ and NELF-B ^30^ are highly conserved in primary sequence among vertebrates, suggesting a conserved contact across species. At the interface-V, the SPT5 KOWx-4 domain packs on Pol II, stabilizes the exit RNA, and contacts the INTS11 (Fig. 2, a, f). As discussed below, the interaction brings RNA to INTS11 for cleavage.

### The Pol II CTD-INTAC interface

The human Pol II CTD consists of 52 consensus heptapeptide repeats (Tyr1-Ser2-Pro3-Thr4-Ser5-Pro6-Ser7) and the phosphorylation levels at Ser2, Ser5, and Ser7 change dynamically throughout the transcription cycle, exhibiting distinct patterns for initiation, elongation, and termination ^32^. Cryo-EM map reveals four putative Pol II CTD segments, indicative of a potential CTD-binding path toward the active center of PP2A-C for dephosphorylation (Fig. 3a, Extended Data Fig. 6, Supplementary Video 2). Pol II and INTAC generate a center-hollowed cradle with the CTD-binding path of INTAC being ~50 Å away from the last modeled RPB1 residue (P1487). This cradle may accommodate Pol II CTD repeats above the three CTD anchoring sites.

**Fig. 3.**
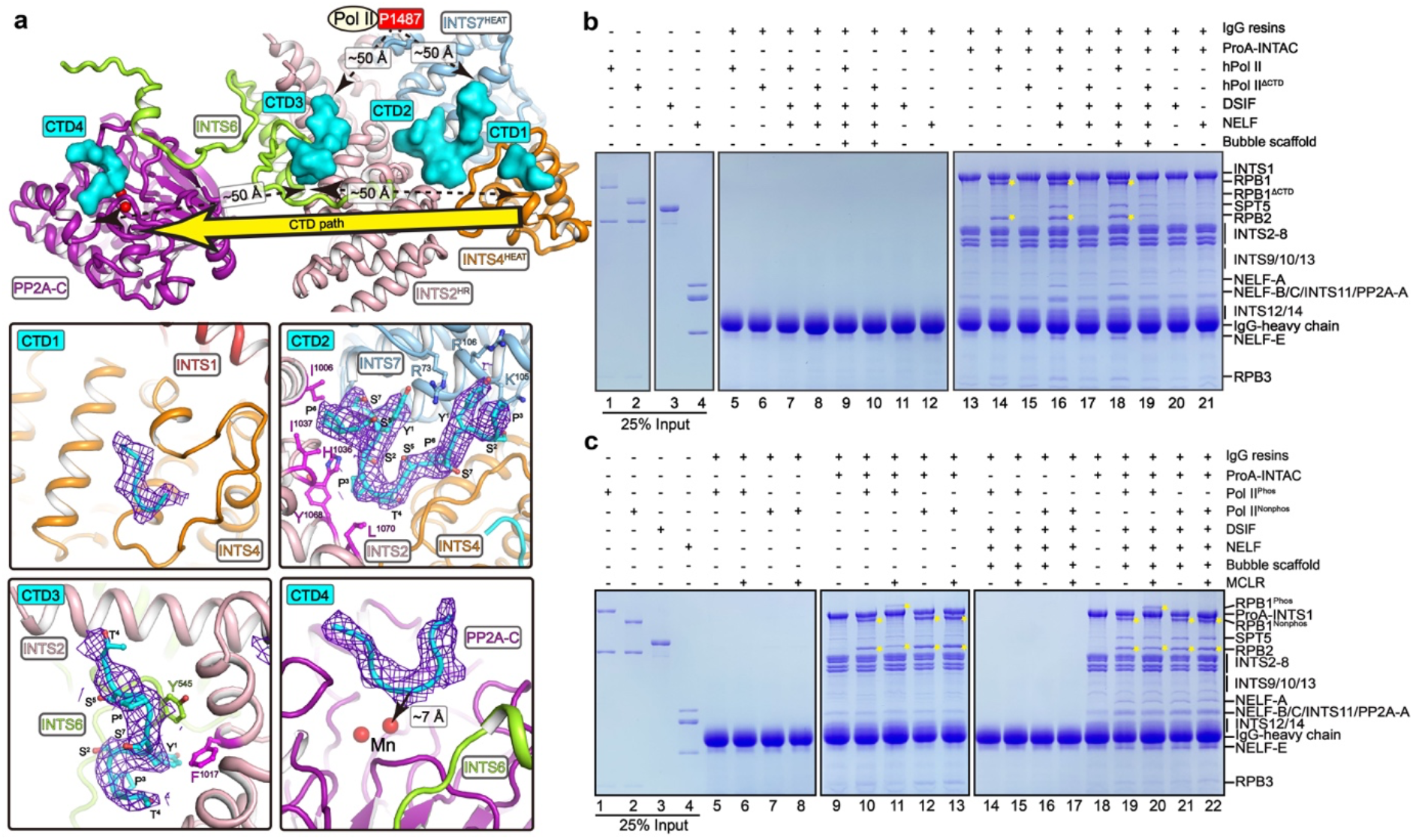
Pol II CTD makes multiple contacts with INTAC and is required for INTAC-Pol II interaction. **a,** Close-up view of Pol II CTD-binding path on INTAC. Four putative CTD fragments are shown in surface and non-relevant subunits/domains were omitted for simplicity. The position of residue P1487 (the last modeled residue) of RPB1 is shown to indicate its distance to CTD fragments. The bottom panels show close-up views of the interactions between CTDs and INTAC with CTDs shown in cryo-EM maps (blue meshes) and structural models. Residues involved in interactions of CTD-2 and CTD-3 are shown in sticks. **b, c,** In vitro pulldown assays using purified INTAC and PEC subcomplexes. Protein A (ProA)-tagged INTAC was incubated with indicated subcomplexes and immobilized onto IgG resins. The unbound proteins were washed and immobilized samples were subjected to SDS-PAGE and Coomassie blue staining. Pol II^phos^ represents pig Pol II that underwent in vitro phosphorylation by TFIIH and hPol II and hPol II^ΔCTD^ represent human Pol II and CTD-truncated human Pol II, respectively. Yellow stars indicate the positions of RPB1 and RPB2, reflecting the binding of Pol II in the reactions.

The CTD-1 to CTD-3 segments span ~50 Å and are sequentially arrayed on the surface of INTAC backbone (Fig. 3a, Extended Data Fig. 6, Supplementary Video 2). The CTD-1 (~5 residues) packs against a relatively hydrophobic pocket of the HEAT repeat of INTS4. The CTD-2 (~13 residues) forms a U-turn coil and packs against the molecular junction of INTS2, INTS4, and INTS7 and is stabilized by a network on interactions. Particularly, two tyrosine (Y^1^) residues anchor on the surface of INTS7 HEAT repeat and sandwich residue R73 of INTS7, generating stacking interaction. The CTD-3 (~8 residues) anchors into a hydrophobic pocket formed by INTS2 and an extending loop of INTS6.

Relatively weak cryo-EM map was observed above the catalytic pocket of PP2A-C (Fig. 3a, Extended Data Fig. 6, c, g). The density is likely derived from Pol II CTD or the N-terminal tail of INTS6 and was termed CTD-4 for simplicity. This U-shaped fragment (~7 residues) is positioned within the substrate-binding groove of PP2A-C. Three central residues are suspended above the catalytic center, in a manner similar to microcystin LR (MCLR, PP2A inhibitor) in the PP2A holoenzyme structure ^33^. The Cα atom of the central residue is ~7 Å away from the near catalytic manganese cation, suggesting a position of phosphorylated Ser5 residue of CTD for dephosphorylation. CTD-4 is ~50 Å away from CTD-3, indicative of a putative CTD path of ~100 Å in length from Pol II body (P1487) to CTD-3 and then to PP2A-C active site.

### Pol II CTD is required for INTAC-PEC interaction

To further investigate how INTAC-PEC complex is assembled, we performed in vitro pulldown assay using immobilized INTAC and individually purified NELF, DSIF (Fig. 3b, Extended Data Fig. 1, c, d). We also overexpressed and purified human Pol II (hPol II) and a CTD-truncated hPol II (hPol II^ΔCTD^) in Expi293F cells (Extended Data Fig. 1a). Consistent with previous studies ^7,19^, INTAC could pull out hPol II with nearly 1:1 stoichiometry (Fig. 3b, lanes 13-14). In contrast, the deletion of CTD impaired INTAC-hPol II interaction (lane 15) and isolated CTD could pull out INTAC (Extended Data Fig. 1c). The immobilized DSIF or NELF exhibited a weak but detectable binding with INTAC (Extended Data Fig. 1d), consistent with previous studies showing their binding to Integrator independent of DNA/RNA ^21,22^. In agreement with this weak interaction and their limited contact with INTAC (Fig. 2), the addition of DSIF and NELF showed nearly undetectable effect on Pol II-INTAC interaction (Fig. 3b, lanes 15, 17). Interestingly, the addition of a DNA-RNA bubble to hPol II^ΔCTD^, along with DSIF and NELF, caused a slight increase in binding to INTAC (lanes 17, 19), suggesting that the exiting RNA may facilitate the binding of PEC to INTAC. The above result underscores the critical role of CTD in the recruitment of INTAC to Pol II and the assembly of INTAC-PEC. NELF, DSIF, and nascent RNA may together facilitate organizing the complex and allow efficient CTD dephosphorylation and RNA cleavage.

The in vitro pulldown assay further showed that INTAC binds and dephosphorylates phosphorylated Pol II (Fig. 3c, lanes 9-10). The addition of PP2A inhibitor hampered the dephosphorylation and binding of Pol II to INTAC (lane 11), as compared to the unphosphorylated Pol II (lanes, 10, 12, 13). The result suggests that CTD phosphorylation partially inhibits INTAC-Pol II interaction and INTAC may tolerate CTD phosphorylation to some extent.

In the presence of NELF and DSIF, INTAC showed a comparable binding to phosphorylated and unphosphorylated Pol II (Fig. 3c, lanes 18-22). NELF and DSIF enhanced INTAC-Pol II interaction and the dephosphorylated CTD may reinforce INTAC-CTD interaction. Structural and biochemical analyses together lead to a model of CTD dephosphorylation. Upon formation of INTAC-PEC complex, un/dephosphorylated CTD occupies the CTD-binding sites of INTAC and evicts, if any, the bound phosphorylated CTD, which is further brought to PP2A-C for dephosphorylation. Thus, INTAC ensures a persistent and complete CTD dephosphorylation for early termination.

### INTS11 is activated upon INTAC-PEC assembly

Consistent with previous studies ^30,31^, the INTAC-PEC structure shows that SPT5 and SPT4, the two DSIF subunits, bind Pol II around stalk, clamp, and wall, with multiple domains surrounding the entry DNA and exit RNA (Figs. 1, 4a, Extended Data Fig. 7, Supplementary Video 3). The SPT5 KOWx-4 and KOW5 domains function as an “RNA clamp” and contact INTAC on INTS9-INTS11 heterodimer. At the interface-V, the INTS11 metallo-β-lactamase domain binds the SPT5 KOWx-4 domain, which bridges the RNA exit tunnel of Pol II and RNA entry tunnel of INTS11. Although DSIF, NELF, and Pol II body are not essentially required for binding to INTAC, INTAC-PEC interactions at interface-I to -V may maintain the overall modular organization and guide RNA to the active center of INTS11 for cleavage.

**Fig. 4.**
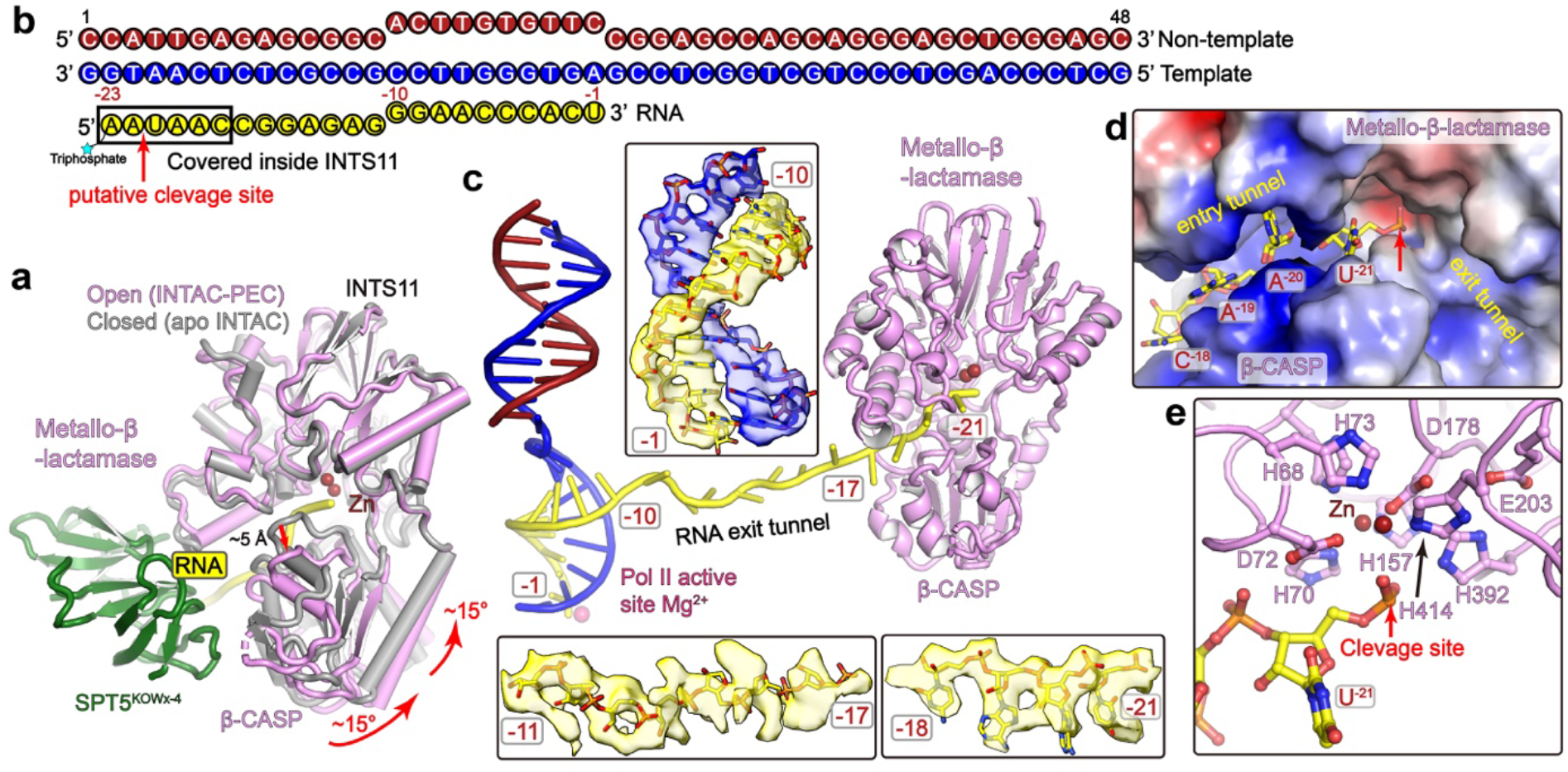
RNA is recognized and cleaved by INTS11 in an active conformation. **a**, Assembly of INTAC-PEC leads to activation of INTS11. The structures of INTS11 in the apo INTAC (grey) ^7^ and INTAC-PEC (pink, this study) are superimposed. Rotation of the β-CASP domain and opening of the RNA entry tunnel are indicated with arrows. The movement of INTS11 likely results from binding of PEC and/or RNA. **b-c**, Schematic diagram (b) and structural model (c) of DNA-RNA bubble. Cryo-EM maps of three parts are shown. Assignment of RNA between −11 to −17 was not accurate and the number of nucleotides was proposed based on the length of extended RNA strand. **d, e,** Recognition of RNA by INTS11 within the RNA entry tunnel (d) and above the active site (e). (d) Electrostatic potential surface of INTS11 is shown and RNA is shown in sticks. (e) Organization of the catalytic center and positioning of RNA for cleavage.

INTS11 exhibits a closed, inactive conformation in the structures of RNA-free INTAC ^7^ and the isolated endonuclease module (INTS4-INTS9-INTS11) ^34^ (Fig. 4a, Extended Data Fig. 7, Supplementary Video 4). Superposition of INTS11 metallo-β-lactamase domain shows that the association of RNA-bound PEC induces a rotation of INTS11 β-CASP domain by ~15 degrees and an opening of the substrate-binding tunnel by ~5 Å, permitting the entry of the RNA for cleavage. Structural comparison suggests an activation of INTS11 upon assembly of INTAC-PEC.

### RNA is brought to INTS11 for cleavage

Among the 23 RNA nucleotides used in structure determination, nucleotides −1 to −10 (relative to the NTP addition site) form DNA-RNA hybrid within Pol II and the following strand (−10 to −16) winds out of RNA exit tunnel (Fig. 4, b, c, Extended Data Fig. 7). A weak but noticeable cryo-EM map showed a linker RNA (−16 to −18), consistent with its lack of protein contact. The four preceding nucleotides (−18 to −21) insert into the RNA entry tunnel of INTS11 with the phosphate and ribose groups well-ordered. The phosphate groups face downward the RNA entry tunnel while the bases face outward and are sandwiched by hydrophobic cleft, consistent with non-RNA sequence specificity of INTS11 (Fig. 4d, Supplementary Video 3).

Nucleotide U^−21^ is suspended near the active site of INTS11 and its phosphate group and the two preceding nucleotides (−22 and −23) were nearly invisible, indicative of a position of RNA cleavage, ~20 nucleotides upstream of the NTP addition site (Fig. 4, c to e, Extended Data Fig. 7b). If the RNA were not cleaved, the phosphate group of U^−21^ would be stabilized by residue H392 of INTS11 with the two phosphate oxygen atoms being ~3 Å away from the two catalytic zinc cations. The preceding nucleotide would be stabilized by residue Y353 of INTS11. The organization of catalytic pocket and the placement of RNA substrate are generally similar to that of cleavage and polyadenylation specificity factor (CPSF) CPSF73 in the histone pre-mRNA cleavage complex (HCC) complex (Extended Data Fig. 7c), which adopts an active state, poised for the cleavage reaction ^35^.

It has been reported that the nucleic acid binding module INTS10-INTS13-INTS14 of INTAC preferentially binds RNA stem loop regions and brings the endonuclease module to target transcripts for cleavage ^36^. This subcomplex was not observed in our structure, possibly due to the lack of stabilization by exposed/uncleaved RNA.

### INTAC generates steric clash with PAF1C and SPT6 of the EC* complex

INTAC has been demonstrated to be involved in NELF- and DSIF-dependent promoter-proximal pausing ^21,22,28^. Recent studies have shown that PAF1 complex (PAF1C) regulates the stabilization of pausing in addition to its role in promoting elongation ^37^. An activated transcription elongation complex EC* is formed by Pol II, PAF1C, DSIF, and SPT6 ^31^. Comparison of the INTAC-PEC structure and the EC* structure shows that the PAF1C subunits PAF1 and LEO1 generate steric hindrance with the tail module of INTAC and CTR9 has clash with INTS2^HR1^ of INTAC ^7^ (Extended Data Fig. 4c). In addition, the SPT6 core around the exit RNA clashes with INTS11 and the SPT6 tSH2 domain has clashes with the INTS6 vWA domain. The above structural comparison shows that PAF1C and INTAC may form independent complexes with paused Pol II, or conformational changes of PAF1C or INTAC are required for their co-existence with a Pol II.

### INTAC works with PEC to orchestrate CTD dephosphorylation and RNA cleavage for premature transcription termination

Mounting evidence has shown that Integrator’s function in snRNA cleavage is rather an exception and that INTAC confers more profound role in transcription regulation. Our study reveals the molecular mechanism by which INTAC shapes the organization of paused Pol II and induces premature transcription termination via the cleavage of nascent RNA by its endonuclease module and the dephosphorylation of Pol II CTD by its phosphatase module. Recruitment of the endonuclease module to nascent RNA requires binding of Pol II CTD to INTAC, which prefers un/dephosphorylated CTD and could tolerate CTD phosphorylation, especially in the presence of NELF and DSIF. The paused Pol II and the cleavage of nascent RNA are thought to destabilize Pol II and lead to exonuclease Xrn2-mediated termination of transcription ^38^. The structure also provides a framework for further study of Integrator’s functions in transcription termination of coding genes and processing of non-coding RNAs ^16–19,26,27^.

During manuscript preparation, Fianu et. al. reported cryo-EM structure of Intergrator-PP2A bound to PEC ^39^. The structure is generally similar to our structure except that RNA was not observed in the RNA entry tunnel of INTS11. Moreover, only one CTD fragment (CTD-2 in our study) was observed, possibly because the Intergrator-PP2A was assembled by mixing three subcomplexes. Nevertheless, our independent study confirms the molecular mechanism of INTAC-PEC assembly and provides additional insights into RNA cleavage by INTS11 and INTAC-mediated Pol II CTD recognition and dephosphorylation.

## Acknowledgments

We thank the Center of Cryo-Electron Microscopy of Fudan University for the supports on data collection. This work was supported by grants from the National key R&D program of China (2016YFA0500700), the National Natural Science Foundation of China (32030055, 31830107, 31821002), the Shanghai Municipal Science and Technology Major Project (2017SHZDZX01), Shanghai Municipal Science and Technology Commission (19JC1411500), the National Ten-Thousand Talent Program (Y. X.), the National Program for support of Top-Notch Young Professionals (Y. X.), and the Strategic Priority Research Program of the Chinese Academy of Sciences (grant no. XDB08000000).

## Author contributions

H. Z. prepared the samples for structural and biochemical analyses with help from F. C., X. W., and X. C; Q. J. collected the data and performed EM analyses and model building with help from Y. Q., W. L., Y. R. and J. C.; Y. X. and H. Z. wrote the manuscript; Y. X. supervised the project.

## Data and materials availability

Cryo-EM maps and atomic coordinates will be deposited in the EMDB and PDB upon the acceptance of this manuscript.

## Methods

### Protein expression and purification

INTAC was overexpressed and purified as previously described ^7^. Pol II was isolated from *S. scrofa thymus* and purified following the reported protocol ^30,31,40,41^. Four residue substitutions (G882S of RBP2, T75I of RPB3, S140N of RPB3, and S126T of RPB6) exist between *S. scrofa* and *H. sapiens* Pol II.

All the purification steps were performed at 4 °C unless otherwise stated. The two full-length open reading frames (ORFs) of human DSIF subunits (SPT4 and SPT5) were separately subcloned into a modified pCAG vector and SPT4 was tagged with an N-terminal 2 × Protein A. Both plasmids were co-transfected to Expi293 cells using PEI (Polysciences) when the cells reached a density of 2.5 × 10^6^/ml. After being cultured at 37 °C for 60 hours, cells were harvested and lysed in lysis buffer containing 50 mM Na-HEPES pH 7.4, 300 mM NaCl, 0.25% CHAPS, 5 mM MgCl_2_, 5 mM adenosine triphosphate (ATP), 10% glycerol (v/v), 2 mM dithiothreitol (DTT), 1 mM phenylmethylsulfonyl fluoride (PMSF), 1 μg/ml aprotinin, 1 μg/ml pepstatin, 1 μg/ml leupeptin for 30 min. The lysate was clarified by centrifugation at 16,000 rotations per minute (rpm) for 30 min with JLA-16.250 rotor (Beckman Coulter), and the supernatant was incubated with immunoglobulin G (IgG) resins (Smart-Lifesciences) overnight. The resins were washed with buffer containing 30 mM Na-HEPES pH 7.4, 300 mM NaCl, 0.1% CHAPS, 2 mM MgCl_2_, 10% glycerol, 2 mM DTT. After on-column cleavage by 3C protease for 4 hours, the immobilized proteins were eluted and further purified by ion exchange chromatography (Mono Q 5/5, GE Healthcare). Peak fractions were assessed by SDS–PAGE followed by Coomassie blue staining. Protein concentration was determined by measuring absorption at 280 nm and using the predicted extinction coefficient for DSIF. Pure fractions were pooled, aliquoted, snap frozen and stored at −80 °C.

NELF was prepared essentially in a similar way as described in DSIF. The four full-length ORFs of human NELF subunits (NELF-A, -B, -D, -E) were separately subcloned into a modified pCAG vector and NELF-E was tagged with an N-terminal 2 × Protein A. The plasmids were co-transfected into Expi293F cells for overexpression. The cells were collected by centrifugation and resuspended in lysis buffer containing 50 mM Na-HEPES pH 7.4, 300 mM NaCl, 0.25% CHAPS, 5 mM MgCl_2_, 5 mM ATP, 10% glycerol (v/v), 2 mM DTT, 1 mM PMSF, 1 μg/ml aprotinin, 1 μg/ml pepstatin, 1 μg/ml leupeptin. After cell lysis, the lysate was cleared by centrifugation and the supernatant was incubated with IgG resins (Smart-Lifesciences) for 4 hours followed by on-column digestion by 3C protease for 4 hours. The eluate was further purified by ion exchange chromatography (Mono Q 5/5, GE Healthcare). Peak fractions were pooled and protein purity was assessed by SDS–PAGE and Coomassie staining. Pure NELF was concentrated and subjected to in vitro dephosphorylation overnight by Lambda Protein Phosphatase (Lambda PP, Beyotime Biotechnology). The dephosphorylated NELF was applied onto a Superdex200 10/300 GL column (GE Healthcare) in a buffer containing 30 mM K-HEPES pH 7.4, 150 mM KCl, 5% glycerol (v/v), 2 mM DTT. Peak fractions containing NELF were pooled, aliquoted, snap frozen, and stored at −80 °C.

### Cryo-EM sample preparation

DNA oligos were purchased from Generay Biotechnology and RNA oligos were purchased from Bioneer. All oligos were resuspended in RNase-free water (200 μM) and stored at −30 °C. The Pol II elongation complex (EC) was assembled on a bubble scaffold with the following nucleic acid sequences as previously reported with minor modifications ^30,31^: template DNA 5′-GCT CCC AGC TCC CTG CTG GCT CCG AGT GGG TTC CGC CGC TCT CAA TGG-3′, non-template DNA 5′-CCA TTG AGA GCG GCA CTT GTG TTC CGG AGC CAG CAG GGA GCT GGG AGC-3′, and RNA 5′-triphosphate-AAU AAC CGG AGA GGG AAC CCA CU-3′. The scaffold contains 10 bp DNA–RNA hybrid, 10-nuclotide bubble, 13 nucleotides of exit RNA, 24 nucleotides of entry DNA and 14 nucleotides of exit DNA. To obtain the DNA-RNA hybrid, template DNA and RNA were mixed with a molar ratio of 1:1.3 and were annealed by incubating the nucleic acids at 95 °C for 10 min and then decreasing the temperature by 1 °C min^−1^ steps to a final temperature of 4 °C in a thermocycler in a buffer containing 20 mM K-HEPES pH 7.4, 60 mM KCl, 3 mM MgCl_2_, and 5% (v/v) glycerol. All concentrations refer to the final concentrations used in complex assembly. To assemble EC, the purified *S. scrofa* Pol II (275 pmol) was incubated with twofold molar excess of the DNA-RNA hybrid for 15 min at 30 °C, shaking at 300 rpm, followed by the addition of twofold molar excess of non-template DNA and further incubation for 15 min at 30 °C. The purified DSIF and NELF were added in a twofold molar excess relative to Pol II for the PEC reconstitution. The sample was incubated for 1 hour at 4 °C, followed by the addition of the purified INTAC (250 pmol) and incubation for another 2 hours at 4 °C. The resulting sample was subjected to gradient fixation (GraFix) ^42^. The glycerol gradient was prepared using light buffer containing 8% (v/v) glycerol, 20 mM K-HEPES pH 7.4, 60 mM KCl, 0.03% CHAPS, 2 3mM MgCl_2_, mM DTT, and heavy buffer containing 40% (v/v) glycerol, 0.0018% glutaraldehyde (Sigma), 20 mM HEPES pH 7.4, 60 mM KCl, 0.03% CHAPS, 3mM MgCl_2_, 2 mM DTT. The centrifugation was performed using an SW60 Ti rotor (Beckman Coulter) at 32,000 rpm at 4°C for 14 hours. Subsequently, peak fractions were pooled and the cross-linking reactions were quenched with 100 mM Tris-HCl pH7.0. The homogeneity of peak fractions was assessed by negative-stain electron microscopy. Fractions of interest were concentrated to about 1.7 mg/ml and dialyzed overnight against a buffer containing 20 mM K-HEPES pH 7.4, 60 mM KCl, 0.8% glycerol, 1 mM tris (2-carboxyethyl) phosphine (TCEP), followed by cryo-EM grid preparation.

For negative-stain EM, 5 μl of freshly purified protein sample was applied onto a glow-discharged copper grid supported by a thin layer of carbon film for 1 min before negative staining by 2% (w/v) uranyl formate at room temperature. The negatively stained grid was loaded onto a FEI Talos L120C microscope operated at 120 kV, equipped with a Ceta CCD camera.

For cryo-EM grid preparation, 4 μl of protein sample (about 0.73 mg/ml) was applied onto a glow-discharged holey carbon grid (Quantifoil Au, R2/2, 300 mesh). After blotting for 3 s, the grid was vitrified by plunging it into liquid ethane using a Vitrobot Mark IV (FEI) operated at 4°C and 100% humidity.

### IgG pulldown assay

Expi293F cells containing overexpressed INTAC complex were pelleted and lysed as previously described ^7^. The supernatant of the cell lysate was incubated with IgG resins for 2 hours at 4°C. The INTAC complex was immobilized on the resins by N-terminal 4×Protein A– tagged INTS1. The resins were extensively washed and resuspended in 450 μl of the binding buffer containing 30 mM K-HEPES pH7.4, 100 mM KCl, 0.1% CHAPS, 3 mM MgCl_2_, 8% glycerol, 2 mM DTT. The purified Pol II or Pol II with deletion of RPB1 CTD (Pol II^CTD^) expressed in Expi293 cells was subjected to removing endogenous RPAP2 by incubating with RPAP2 antibody (Abclonal) on Protein G resins. The resulting Pol II, Pol II^ΔCTD^, or their the mixture with DSIF and NELF, in the presence or absence of a bubble scaffold was individually incubated with INTAC-immobilized IgG resins for 2 hours at 4°C. The resins were extensively washed with the binding buffer, and the bound proteins were subjected to SDS-PAGE followed by Coomassie blue staining. Other IgG pulldown assays were performed in a similar approach as described above.

### Cryo-EM data collection and image processing

Cryo-EM data were collected on a Titan Krios electron microscope (FEI) operated at 300 kV at the Cryo-EM platform of Fudan University, equipped with a K2 summit direct detector (Gatan) and a GIF quantum energy filter (Gatan) set to a slit width of 20 eV. Automated data acquisition was carried out with Serial EM software in the superresolution mode ^43^ at a nominal magnification 130,000×, corresponding to a calibrated pixel size of 1.054 Å, and a defocus range from −1.5 to −2.5 μm. Each image stack was dose fractionated to 32 frames with a total exposure dose of about 50 e−/Å2 and exposure time of 6.72 s. The image stacks were motion-corrected and dose-weighted using MotionCorr2 ^44^. The contrast transfer function (CTF) parameters were estimated by CTFFIND-4.1 from non-dose weighted micrographs. About 51,000 particles autopicked from 2000 micrographs were subjected into two-dimensional (2D) classification in RELION v3.0 ^45^ and ab initio reconstruction by cryoSPARC v2 ^46^. The 3D initial model was low-passed and used as references for subsequent particle-picking and 3D classification. The following procedures of image processing were performed using RELION for dose-weighted micrographs, 1,237,927 particles were autopicked from 13,956 micrographs for further data processing. After several rounds of 3D classification, 47,736 good particles were selected for further no-alignment 3D classification. Because of the relatively flexible organization between INTAC and PEC, the mask of INTS1-INTS6-INTS9-INTS11-PEC was applied to no-alignment 3D classification to separate the weakly associated INTAC-PEC. Finally, 21,304 particles (stably associated INTAC-PEC) were subjected to 3D-autorefinement, postprocessing, CTF refinement and Bayesian polishing , yielding a reconstruction of INTAC-PEC at 4.47 Å resolution. In order to improve the map quality for model building, focused classification and refinement were used. Afterwards, selected particles were postprocessed, CTF-refined, Bayesian polished and generated reconstructions of the INTAC at 3.92 Å (78,201 particles) , INTS2-INTS7-CTD at 3.78 Å (78,201 particles), PP2A-AC at 4.10 Å (70,374 particles), INTS9-INTS11-Pol II-DSIF at 3.99 Å (42,152 particles), INTS9-INTS11-RNA at 4.10 Å (38,826 particles) and INTS11-RNA at 3.99 Å (38,826 particles). The reported resolutions above are based on the gold-standard Fourier shell correlation (FSC) 0.143 criterion. All the visualization and evaluation of 3D density maps were performed with UCSF Chimera 47 or UCSF ChimeraX ^48^, and the local resolution variations were calculated using ResMap. The above procedures of data processing are summarized in Extended Data Fig. 2.

### Model building and structure refinement

The structural model of INTAC-PEC was built according to the 4.47 Å INTAC-PEC cryo-EM map and corresponding focused refined maps. The structures of human INTAC (PDB: 7CUN) and PEC (PDB: 6GML) were used to guide modelling of INTAC-PEC, which were docked into the INTAC-PEC cryo-EM map by rigid body fitting using UCSF Chimera ^47^ and were manually adjusted using COOT ^49^. The models of INTS1, INTS2, INTS4, INTS8, INTS9 and INTS11 were further optimized in the guidance of the protein structures predicted by AlphaFold ^50^. To build the model of INTS11 (active conformation) and RNA (−18 to −21), the homologous structure of CPSF with nascent RNA (PDB: 6V4X) was used as a reference according to the INTS9-INTS11-RNA map and the model of RNA (−1 to −17) was built using map INTS11-RNA. The structural model of the INTAC-PEC complex was refined against the 4.47 Å overall map in real space with PHENIX ^51^ and validated through examination of Ramachandran plot statistics, a MolProbity score ^52^, and a EMRinger score ^53^. The statistics of the map reconstruction and model refinement are summarized in Extended Data Table 1. Each focused refined maps were used to create the composite map using UCSF ChimeraX ^48^. The composite map was used in Fig. 1a and Supplementary Video 1. Map and model representations in the figures and videos were prepared by PyMOL and UCSF ChimeraX ^48^.

## Reference

1 Jonkers, I., Kwak, H. & Lis, J. T. Genome-wide dynamics of Pol II elongation and its interplay with promoter proximal pausing, chromatin, and exons. Elife 3, e02407, doi:10.7554/eLife.02407 (2014).

2 Day, D. S. et al. Comprehensive analysis of promoter-proximal RNA polymerase II pausing across mammalian cell types. Genome Biol 17, 120, doi:10.1186/s13059-016-0984-2 (2016).

3 Core, L. & Adelman, K. Promoter-proximal pausing of RNA polymerase II: a nexus of gene regulation. Genes Dev 33, 960–982, doi:10.1101/gad.325142.119 (2019).

4 Jonkers, I. & Lis, J. T. Getting up to speed with transcription elongation by RNA polymerase II. Nat Rev Mol Cell Biol 16, 167–177, doi:10.1038/nrm3953 (2015).

5 Chen, F. X., Smith, E. R. & Shilatifard, A. Born to run: control of transcription elongation by RNA polymerase II. Nat Rev Mol Cell Biol 19, 464–478, doi:10.1038/s41580-018-0010-5 (2018).

6 Kamieniarz-Gdula, K. & Proudfoot, N. J. Transcriptional Control by Premature Termination: A Forgotten Mechanism. Trends Genet 35, 553–564, doi:10.1016/j.tig.2019.05.005 (2019).

7 Zheng, H. et al. Identification of Integrator-PP2A complex (INTAC), an RNA polymerase II phosphatase. Science 370, eabb5872, doi:10.1126/science.abb5872 (2020).

8 Wu, C. H. et al. NELF and DSIF cause promoter proximal pausing on the hsp70 promoter in Drosophila. Genes Dev 17, 1402–1414, doi:10.1101/gad.1091403 (2003).

9 Missra, A. & Gilmour, D. S. Interactions between DSIF (DRB sensitivity inducing factor), NELF (negative elongation factor), and the Drosophila RNA polymerase II transcription elongation complex. Proc Natl Acad Sci U S A 107, 11301–11306, doi:10.1073/pnas.1000681107 (2010).

10 Wada, T. et al. DSIF, a novel transcription elongation factor that regulates RNA polymerase II processivity, is composed of human Spt4 and Spt5 homologs. Genes Dev 12, 343–356, doi:10.1101/gad.12.3.343 (1998).

11 Yamaguchi, Y. et al. NELF, a multisubunit complex containing RD, cooperates with DSIF to repress RNA polymerase II elongation. Cell 97, 41–51, doi:10.1016/s0092-8674(00)80713-8 (1999).

12 Evans, R., Weber, J., Ziff, E. & Darnell, J. E. Premature termination during adenovirus transcription. Nature 278, 367–370, doi:10.1038/278367a0 (1979).

13 Mendoza-Figueroa, M. S., Tatomer, D. C. & Wilusz, J. E. The Integrator Complex in Transcription and Development. Trends Biochem Sci 45, 923–934, doi:10.1016/j.tibs.2020.07.004 (2020).

14 Baillat, D. & Wagner, E. J. Integrator: surprisingly diverse functions in gene expression. Trends Biochem Sci 40, 257–264, doi:10.1016/j.tibs.2015.03.005 (2015).

15 Cazalla, D., Xie, M. & Steitz, J. A. A primate herpesvirus uses the integrator complex to generate viral microRNAs. Mol Cell 43, 982–992, doi:10.1016/j.molcel.2011.07.025 (2011).

16 Xie, M. et al. The host Integrator complex acts in transcription-independent maturation of herpesvirus microRNA 3’ ends. Genes Dev 29, 1552–1564, doi:10.1101/gad.266973.115 (2015).

17 Lai, F., Gardini, A., Zhang, A. & Shiekhattar, R. Integrator mediates the biogenesis of enhancer RNAs. Nature 525, 399–403, doi:10.1038/nature14906 (2015).

18 Skaar, J. R. et al. The Integrator complex controls the termination of transcription at diverse classes of gene targets. Cell Res 25, 288–305, doi:10.1038/cr.2015.19 (2015).

19 Baillat, D. et al. Integrator, a multiprotein mediator of small nuclear RNA processing, associates with the C-terminal repeat of RNA polymerase II. Cell 123, 265–276, doi:10.1016/j.cell.2005.08.019 (2005).

20 Gardini, A. et al. Integrator regulates transcriptional initiation and pause release following activation. Mol Cell 56, 128–139, doi:10.1016/j.molcel.2014.08.004 (2014).

21 Yamamoto, J. et al. DSIF and NELF interact with Integrator to specify the correct post-transcriptional fate of snRNA genes. Nat Commun 5, 4263, doi:10.1038/ncomms5263 (2014).

22 Stadelmayer, B. et al. Integrator complex regulates NELF-mediated RNA polymerase II pause/release and processivity at coding genes. Nat Commun 5, 5531, doi:10.1038/ncomms6531 (2014).

23 Lykke-Andersen, S. et al. Integrator is a genome-wide attenuator of non-productive transcription. Mol Cell 81, 514–529 e516, doi:10.1016/j.molcel.2020.12.014 (2021).

24 Huang, K. L. et al. Integrator Recruits Protein Phosphatase 2A to Prevent Pause Release and Facilitate Transcription Termination. Mol Cell 80, 345–358 e349, doi:10.1016/j.molcel.2020.08.016 (2020).

25 Beckedorff, F. et al. The Human Integrator Complex Facilitates Transcriptional Elongation by Endonucleolytic Cleavage of Nascent Transcripts. Cell Rep 32, 107917, doi:10.1016/j.celrep.2020.107917 (2020).

26 Tatomer, D. C. et al. The Integrator complex cleaves nascent mRNAs to attenuate transcription. Genes Dev 33, 1525–1538, doi:10.1101/gad.330167.119 (2019).

27 Rubtsova, M. P. et al. Integrator is a key component of human telomerase RNA biogenesis. Sci Rep 9, 1701, doi:10.1038/s41598-018-38297-6 (2019).

28 Elrod, N. D. et al. The Integrator Complex Attenuates Promoter-Proximal Transcription at Protein-Coding Genes. Mol Cell 76, 738–752 e737, doi:10.1016/j.molcel.2019.10.034 (2019).

29 Vervoort, S. J. et al. The PP2A-Integrator-CDK9 axis fine-tunes transcription and can be targeted therapeutically in cancer. Cell 184, 3143–3162 e3132, doi:10.1016/j.cell.2021.04.022 (2021).

30 Vos, S. M., Farnung, L., Urlaub, H. & Cramer, P. Structure of paused transcription complex Pol II-DSIF-NELF. Nature 560, 601–606, doi:10.1038/s41586-018-0442-2 (2018).

31 Vos, S. M. et al. Structure of activated transcription complex Pol II-DSIF-PAF-SPT6. Nature 560, 607–612, doi:10.1038/s41586-018-0440-4 (2018).

32 Harlen, K. M. & Churchman, L. S. The code and beyond: transcription regulation by the RNA polymerase II carboxy-terminal domain. Nat Rev Mol Cell Biol 18, 263–273, doi:10.1038/nrm.2017.10 (2017).

33 Xu, Y. et al. Structure of the protein phosphatase 2A holoenzyme. Cell 127, 1239–1251, doi:10.1016/j.cell.2006.11.033 (2006).

34 Pfleiderer, M. M. & Galej, W. P. Structure of the catalytic core of the Integrator complex. Mol Cell 81, 1246–1259 e1248, doi:10.1016/j.molcel.2021.01.005 (2021).

35 Sun, Y. et al. Structure of an active human histone pre-mRNA 3’-end processing machinery. Science 367, 700–703, doi:10.1126/science.aaz7758 (2020).

36 Sabath, K. et al. INTS10-INTS13-INTS14 form a functional module of Integrator that binds nucleic acids and the cleavage module. Nat Commun 11, 3422, doi:10.1038/s41467-020-17232-2 (2020).

37 Chen, F. X. et al. PAF1, a Molecular Regulator of Promoter-Proximal Pausing by RNA Polymerase II. Cell 162, 1003–1015, doi:10.1016/j.cell.2015.07.042 (2015).

38 Proudfoot, N. J. Transcriptional termination in mammals: Stopping the RNA polymerase II juggernaut. Science 352, aad9926, doi:10.1126/science.aad9926 (2016).

39 Fianu, I. et al. Structural basis of Integrator-mediated transcription regulation. Science 374, 883–887, doi:10.1126/science.abk0154 (2021).

40 Chen, X. et al. Structures of the human Mediator and Mediator-bound preinitiation complex. Science 372, eabg0635, doi:10.1126/science.abg0635 (2021).

41 Chen, X. et al. Structural insights into preinitiation complex assembly on core promoters. Science 372, eaba8490, doi:10.1126/science.aba8490 (2021).

42 Kastner, B. et al. GraFix: sample preparation for single-particle electron cryomicroscopy. Nat Methods 5, 53–55, doi:10.1038/nmeth1139 (2008).

43 Mastronarde, D. N. Automated electron microscope tomography using robust prediction of specimen movements. J Struct Biol 152, 36–51, doi:10.1016/j.jsb.2005.07.007 (2005).

44 Zheng, S. Q. et al. MotionCor2: anisotropic correction of beam-induced motion for improved cryo-electron microscopy. Nat Methods 14, 331–332, doi:10.1038/nmeth.4193 (2017).

45 Scheres, S. H. RELION: implementation of a Bayesian approach to cryo-EM structure determination. J Struct Biol 180, 519–530, doi:10.1016/j.jsb.2012.09.006 (2012).

46 Punjani, A., Rubinstein, J. L., Fleet, D. J. & Brubaker, M. A. cryoSPARC: algorithms for rapid unsupervised cryo-EM structure determination. Nat Methods 14, 290–296, doi:10.1038/nmeth.4169 (2017).

47 Pettersen, E. F. et al. UCSF Chimera--a visualization system for exploratory research and analysis. J Comput Chem 25, 1605–1612, doi:10.1002/jcc.20084 (2004).

48 Pettersen, E. F. et al. UCSF ChimeraX: Structure visualization for researchers, educators, and developers. Protein Sci 30, 70–82, doi:10.1002/pro.3943 (2021).

49 Emsley, P. & Cowtan, K. Coot: model-building tools for molecular graphics. Acta Crystallogr D Biol Crystallogr 60, 2126–2132, doi:10.1107/S0907444904019158 (2004).

50 Jumper, J. et al. Highly accurate protein structure prediction with AlphaFold. Nature 596, 583–589, doi:10.1038/s41586-021-03819-2 (2021).

51 Adams, P. D. et al. PHENIX: building new software for automated crystallographic structure determination. Acta Crystallogr D Biol Crystallogr 58, 1948–1954, doi:10.1107/s0907444902016657 (2002).

52 Chen, V. B. et al. MolProbity: all-atom structure validation for macromolecular crystallography. Acta Crystallogr D Biol Crystallogr 66, 12–21, doi:10.1107/S0907444909042073 (2010).

53 Barad, B. A. et al. EMRinger: side chain–directed model and map validation for 3D cryo-electron microscopy. Nature Methods 12, 943–946, doi:10.1038/nmeth.3541 (2015).

