## Supplemental Figures for "Structural basis of INTAC-regulated transcription"

### Extended data figures and tables

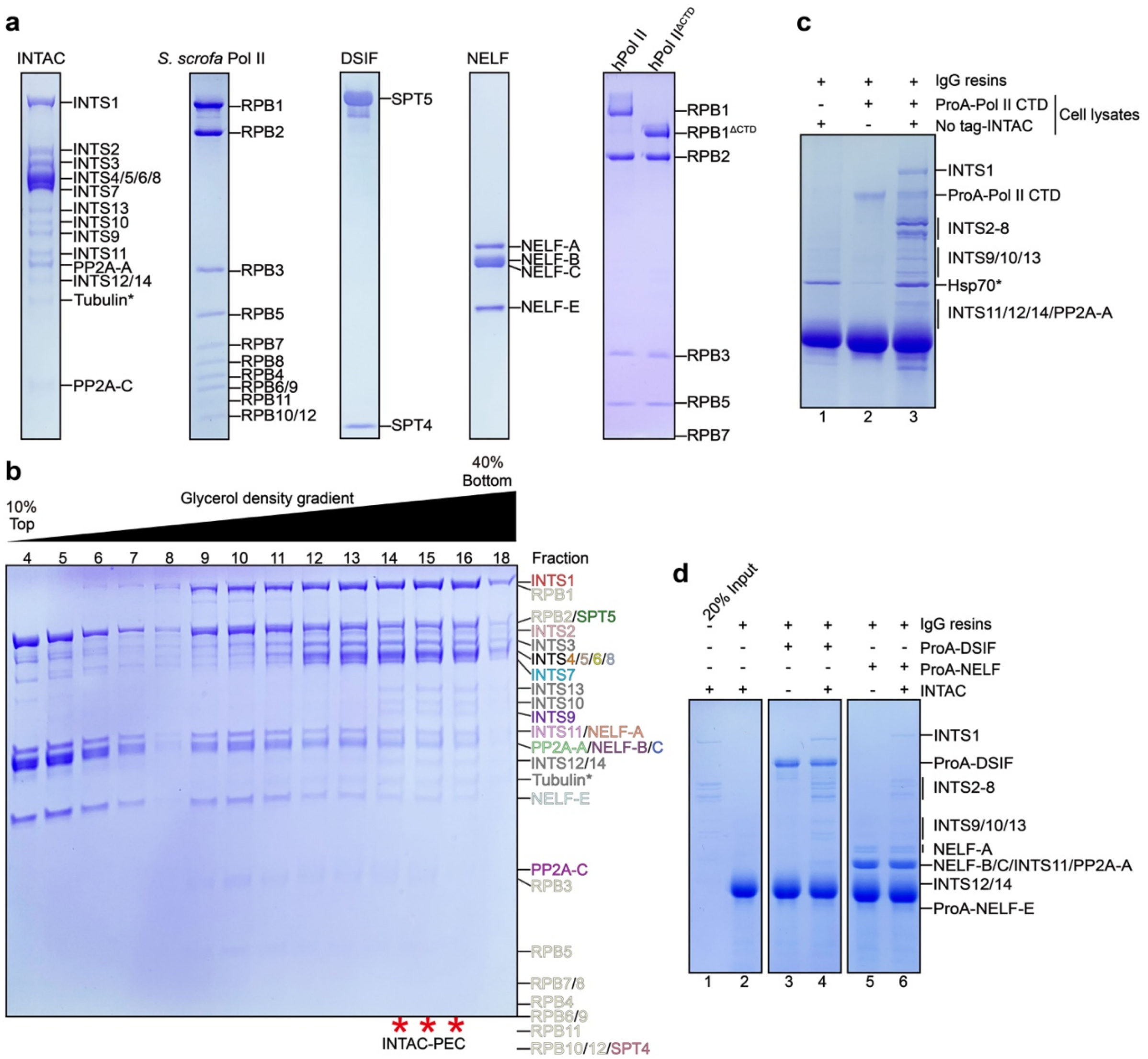

**Extended Data Fig. 1 | Protein purification and complex assembly.**

**a,** Protein complexes used in biochemical and structural analyses. The samples were subjected to SDS-PAGE and stained using Coomassie blue. Pol II represents RNA polymerase II purified from S. scrofa and hPol II represents the Pol II overexpressed and purified from Expi293F cells. hPol IIΔCTD represents hPol II containing truncation of RPB1 CTD (residues 1593-1970). To better reveal phosphorylated hPol II and CTD truncation of RPB1, the lower bands of Pol II subunits were omitted. **b,** Glycerol density gradient of the mixture of INTAC and PEC. Peak fractions of the assembled INTAC-PEC complex are indicated with red stars below. **c-d,** IgG pulldown assay using protein A (ProA)-tagged CTD (c) and DSIF and NELF (d) to test their binding of INTAC. Contaminating proteins (Hsp70 and Tubulin) are indicated with black stars.

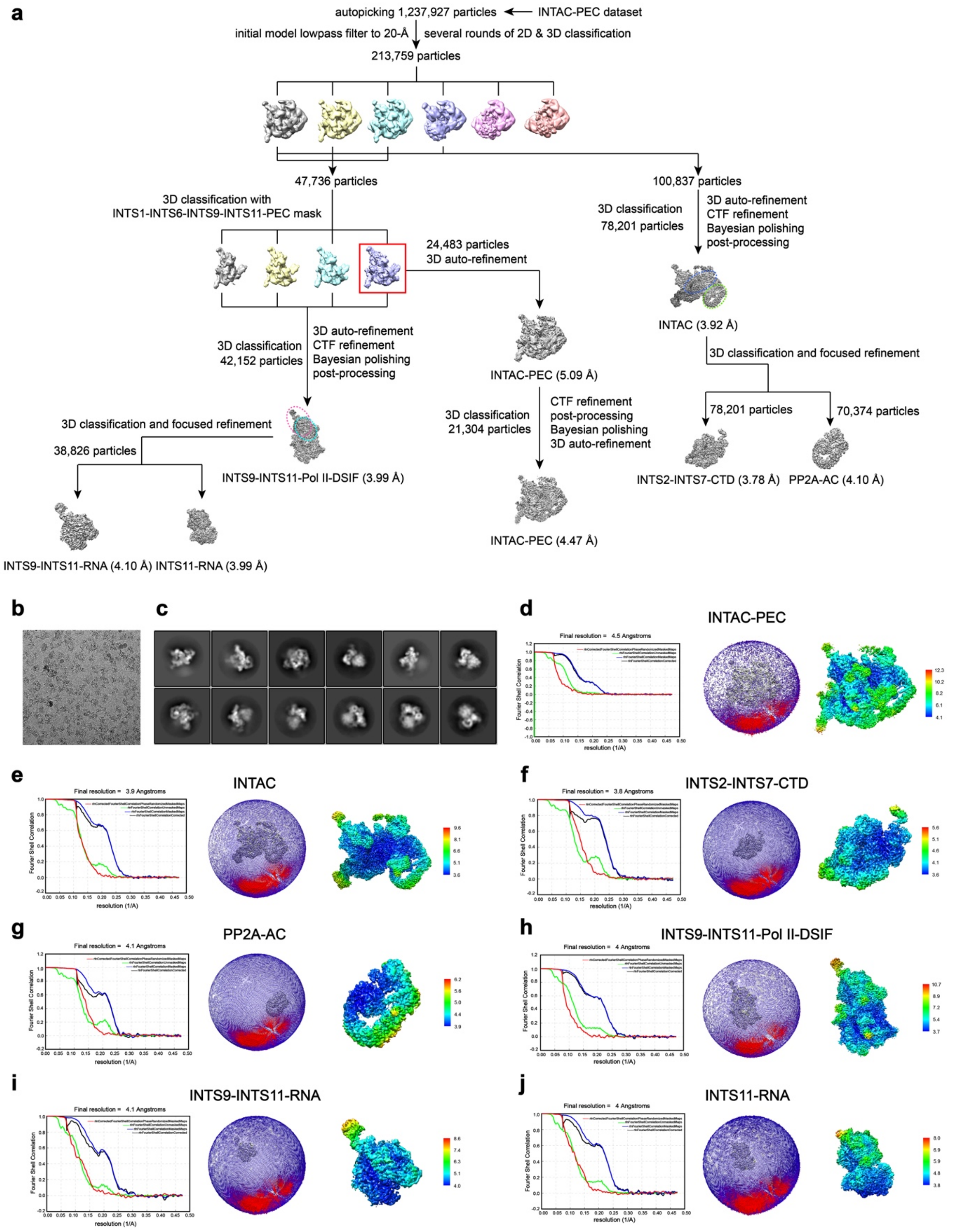

**Extended Data Fig. 2 | Data processing.**

**a,** Flow-charts of the cryo-EM image processing and 3D reconstructions. **b-c,** Representative cryo-EM raw micrograph (b) and 2D classification (c) of INTAC-PEC. **d-j,** The GSFSC curves, angular distribution plots, and local resolution estimation of the cryo-EM maps of global refinement (d) and focused refinements (e-j).

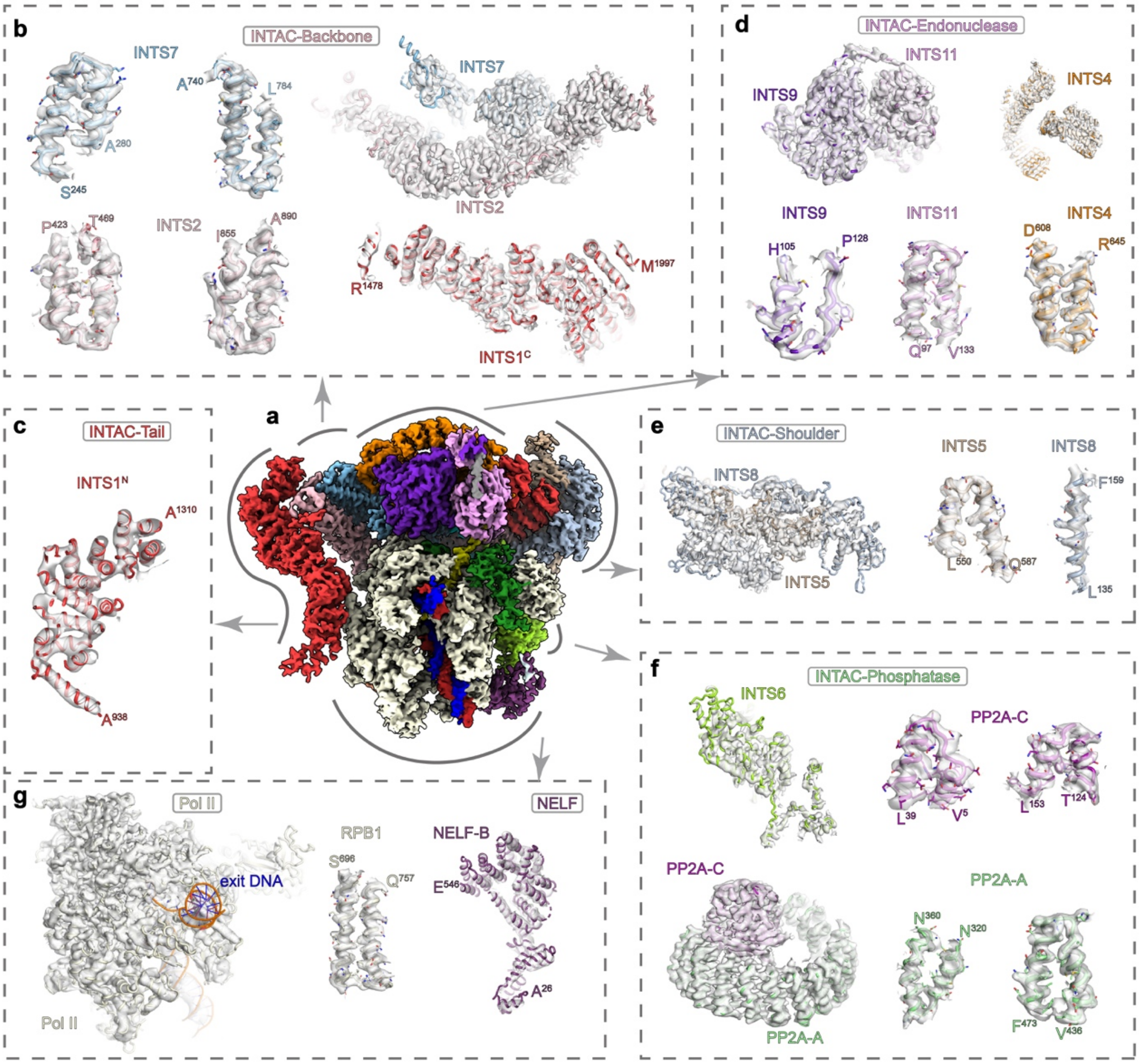

**Extended Data Fig. 3 | Cryo-EM maps and structural models.**

**a,** Composite cryo-EM map of INTAC-PEC shown as in Fig. 1. The positions of each submodule shown in (**b-g**) are indicated on the overall map. **b-g,** Cryo-EM density for each submodule, and close-up views of representative structural models with the corresponding cryo-EM maps shown in surface. Proteins are shown in ribbon and sticks (side chains). Most of the side chains fit into the cryo-EM map, indicating the model was built correctly.

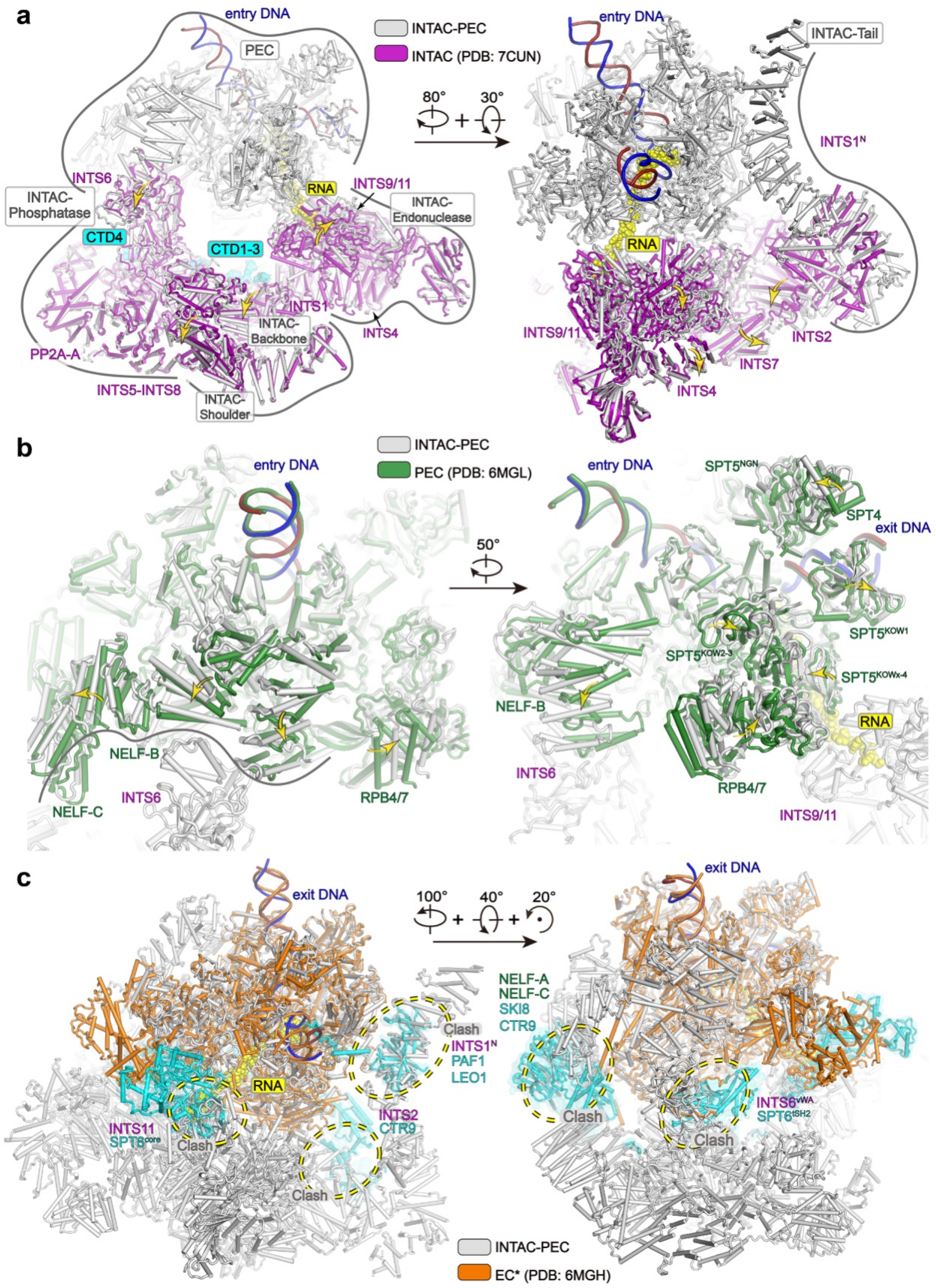

**Extended Data Fig. 4 | Conformational changes of INTAC and PEC upon formation of INTAC-PEC and steric clashes between INTAC and EC* modules.**

Structural comparison of INTAC-PEC with INTAC (PDB: 7CUN) [1] (a), PEC (PDB: 6MGL) [2] (b), and EC* [3] (c). For clarity, INTAC-PEC is colored in grey and other complexes are colored as indicated. Structural differences are highlighted and modular displacements are indicated with arrows. Putative steric clashes are indicated with dashed circles in (c). Two different views are shown for each comparison.

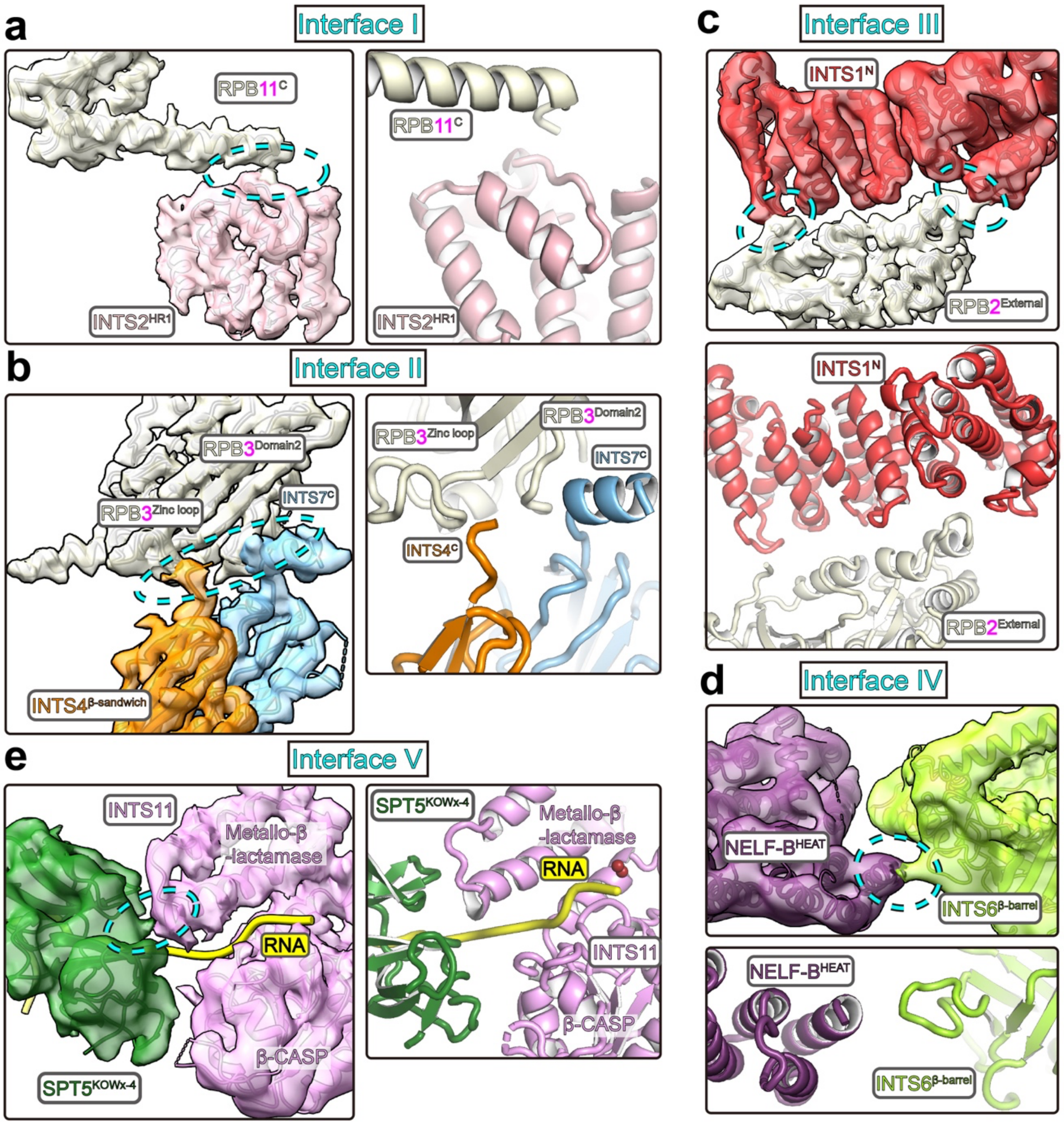

**Extended Data Fig. 5 | Interfaces between INTAC and PEC.**

**a-e,** Interface-I to -V are shows with structural models (right or bottom panels) and structural models covered by transparent cryo-EM maps (left or upper panels) as shown in Fig. 2. The two panels are shown in similar orientation. Contacts are highlighted with dashed circles. Interactions between INTAC and Pol II CTD are omitted here and shown in Extended Data Fig. 6.

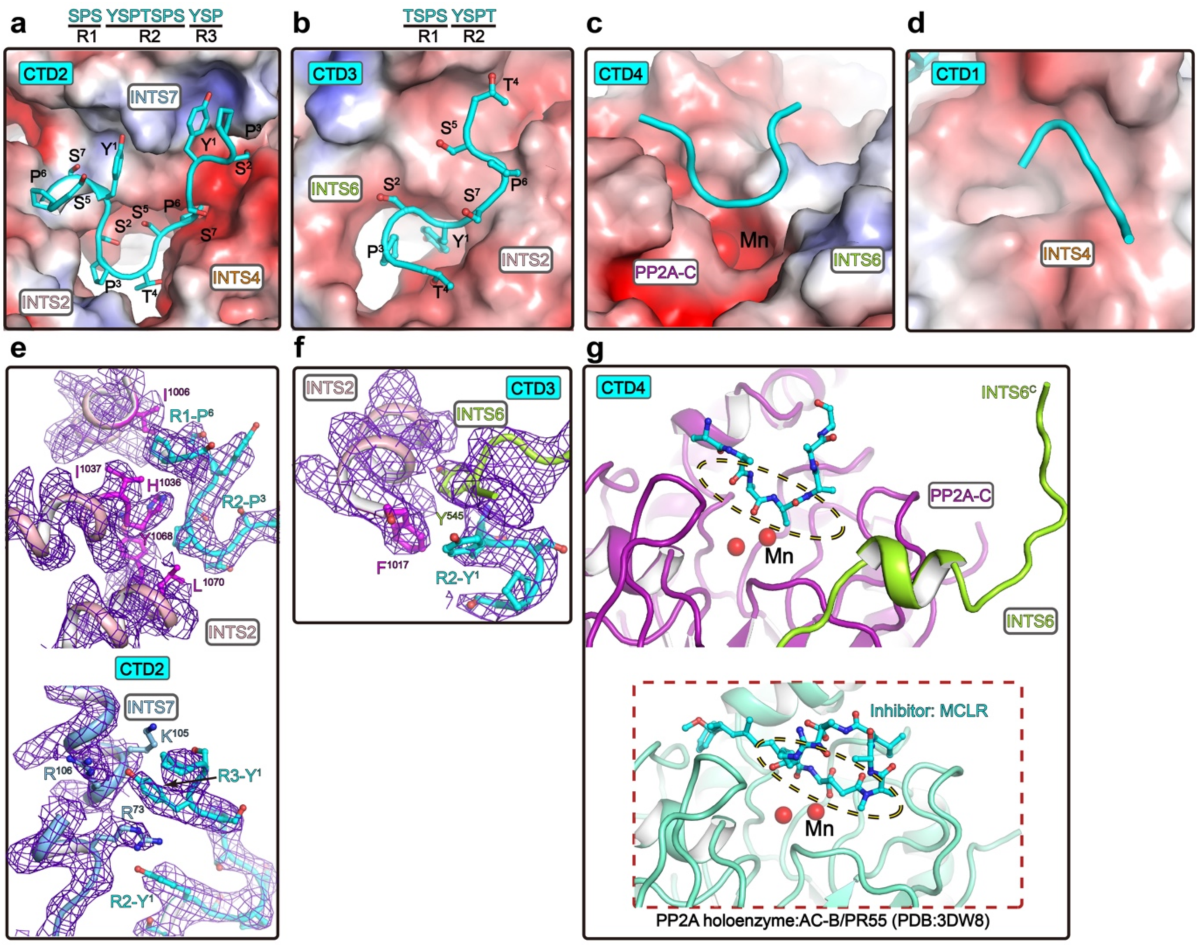

**Extended Data Fig. 6 | Binding of Pol II CTD to INTAC backbone and PP2A-C.**

**a-d,** Interactions between INTAC and four CTD segments. Electrostatic potential surface of INTAC is shown. CTD2 and CTD3 are shown in sticks with hated peptide indicated. As shown Fig. 3, the cryo-EM maps did not support unambiguous assignment of CTD1 and CTD4 CTD. The two segments are shown in cartoon. **e-f,** Cryo-EM maps around CTD2 and CTD3 segments and CTD-binding site of INTAC. Cryo-EM maps are shown in blue meshes. Residues that potentially involved in recognition of CTD are shown in sticks. **g,** Comparison of CTD4 and MCLR (PP2A inhibitor) in INTAC-PEC and PP2A holoenzyme [4] structures. The two PP2A-C are shown in a similar orientation.

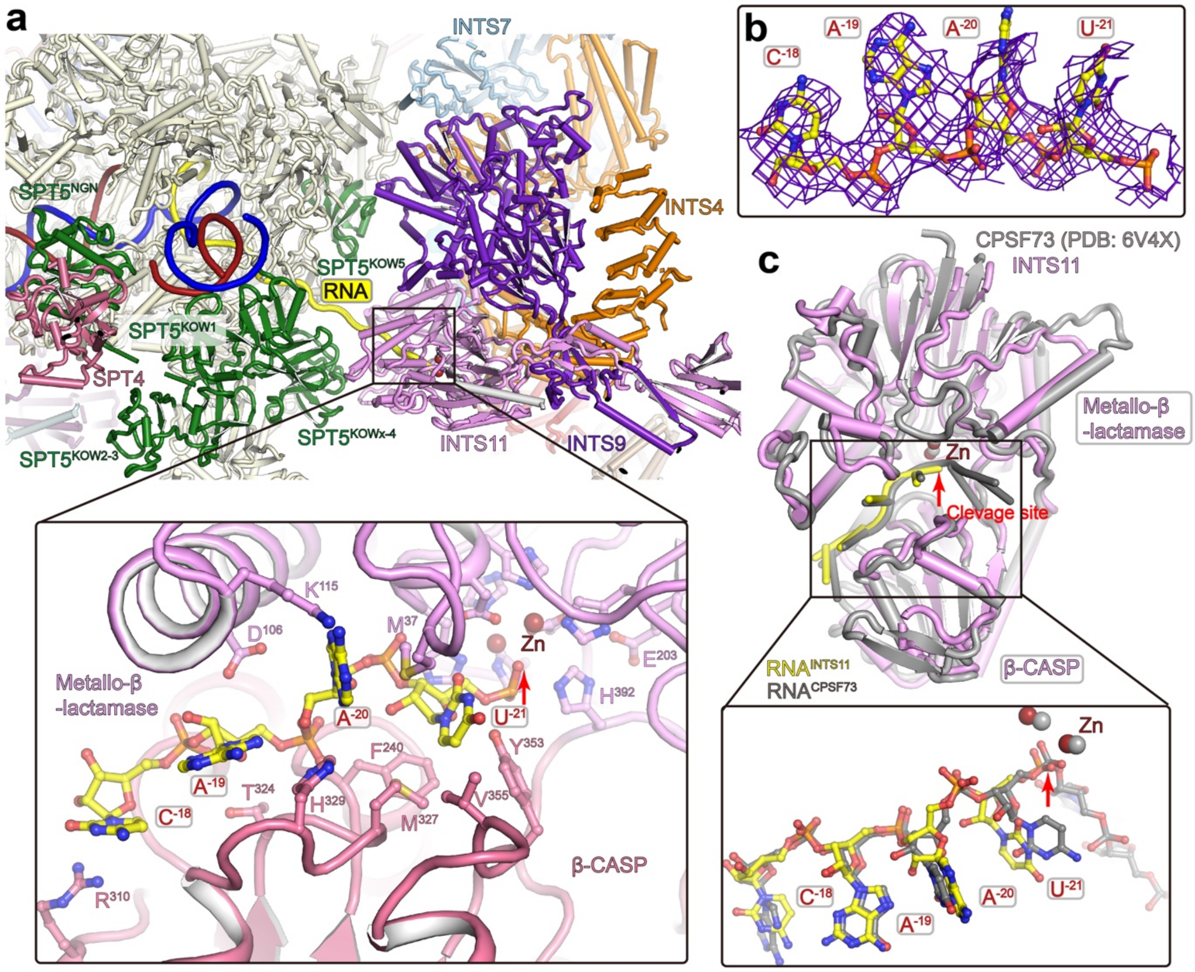

**Extended Data Fig. 7 | Binding of RNA within the RNA entry tunnel of INTS11.**

**a,** Contact between PEC and endonuclease module of INTAC. Close-up view of RNA within the RNA entry tunnel of INTS11 is shown below. **b,** Cryo-EM map of the RNA within INTS11 is shown in mesh and the RNA is shown in sticks. **c,** Structural comparison of the RNA-bound INTS11 in INTAC-PEC and the RNA-bound CPSF73 in HCC (PDB: 6V4X) [5]. Note that RNA was not cleaved in HCC complex.

**Extended Data Table 1. Cryo-EM data collection, refinement and validation statistics**

| | #1 INTAC-PEC (EMDB-xxxx) (PDB xxxx) | #2 INTS2-INTS7-CTD | #3 INTS9-INTS11-RNA |
|---|---|---|---|
| **Data collection and processing** | | | |
| Magnification | 130, 000x | 130, 000x | 130, 000x |
| Voltage (kV) | 300 | 300 | 300 |
| Electron exposure (e–/Å$^2$) | 50 | 50 | 50 |
| Defocus range (μm) | -1.5 to -2.5 | -1.5 to -2.5 | -1.5 to -2.5 |
| Pixel size (Å) | 1.054 | 1.054 | 1.054 |
| Symmetry imposed | C1 | C1 | C1 |
| Initial particle images (no.) | 1, 237, 927 | 1, 237, 927 | 1, 237, 927 |
| Final particle images (no.) | 21, 304 | 78,201 | 38,826 |
| Map resolution (Å) | 4.5 | 3.8 | 4.1 |
| FSC threshold | 0.143 | 0.143 | 0.143 |
| Map resolution range (Å) | 4.1-12.3 | 3.6-5.6 | 4.0-8.6 |
| **Refinement** | | | |
| Initial model used (PDB code) | 7CUN, 6GML | | |
| Model resolution (Å) | 4.3 | | |
| FSC threshold | 0.5 | | |
| Map sharpening *B* factor (Å$^2$) | -96 | -90 | -139 |
| Model composition | | | |
| Non-hydrogen atoms | 113, 053 | | |
| Protein residues | 14, 742 | | |
| Nucleotide residues | 102 | | |
| Ligands | MN: 2, MG: 1, ZN: 11 | | |
| *B* factors (Å$^2$) | | | |
| Protein | 125.16 | | |
| Nucleotide | 294.90 | | |
| Ligand | 197.15 | | |
| R.m.s. deviations | | | |
| Bond lengths (Å) | 0.007 | | |
| Bond angles (°) | 0.859 | | |
| Validation | | | |
| MolProbity score | 2.17 | | |
| Clashscore | 20.04 | | |
| Poor rotamers (%) | 0.00 | | |
| Ramachandran plot | | | |
| Favored (%) | 94.61 | | |
| Allowed (%) | 5.23 | | |

| | |
|---|---|
| Disallowed (%) | 0.16 |

**Supplementary Video 1**

Composite cryo-EM map and structural model of INTAC-PEC.

**Supplementary Video 2**

The CTD-binding path on INTAC. Cryo-EM map and structural model of CTD-binding path and four putative CTD fragments are shown. Pol II is shown to indicate its relative position to the CTD-binding path. The last modeled residue of RPB1 (P1487) is indicated with red ball.

**Supplementary Video 3**

Cryo-EM map and structural model of PEC and endonuclease module of INTAC. The RNA is colored in yellow.

**Supplementary Video 4**

Binding of RNA-bound PEC to INTAC leads to activation of INTS11. Non-related modules were omitted for simplicity.

**References**

1 Zheng, H. *et al.* Identification of Integrator-PP2A complex (INTAC), an RNA polymerase II phosphatase. *Science* **370**, eabb5872, doi:10.1126/science.abb5872 (2020).

2 Vos, S. M., Farnung, L., Urlaub, H. & Cramer, P. Structure of paused transcription complex Pol II-DSIF-NELF. *Nature* **560**, 601-606, doi:10.1038/s41586-018-0442-2 (2018).

3 Vos, S. M. *et al.* Structure of activated transcription complex Pol II-DSIF-PAF-SPT6. *Nature* **560**, 607-612, doi:10.1038/s41586-018-0440-4 (2018).

4 Xu, Y. *et al.* Structure of the protein phosphatase 2A holoenzyme. *Cell* **127**, 1239-1251, doi:10.1016/j.cell.2006.11.033 (2006).

5 Sun, Y. *et al.* Structure of an active human histone pre-mRNA 3'-end processing machinery. *Science* **367**, 700-703, doi:10.1126/science.aaz7758 (2020).
